# High-quality genome assemblies of 152 root commensal bacteria from the model legume *Lotus japonicus*

**DOI:** 10.1101/2025.01.02.630998

**Authors:** Adrián Gómez-Repollés, Eber Villa-Rodriguez, Shaun Ferguson, Simona Radutoiu

## Abstract

Bacterial culture collections represent a valuable tool for mechanistic understanding of microbiome assemblies and are increasingly used to assemble tailored synthetic communities to characterize their microbe-microbe interactions and those with the environment. Given the size of these collections, short-read sequencing is primarily used to capture the encoded genetic information. Whilst sufficient for many microbiome studies, this approach is not amenable for understanding bacterial genome evolution or detailed genetic analyses at the entire genome level. Here we report the assembly of 152 full bacterial genomes from the *Lj-*SPHERE, the *Lotus japonicus* collection of root commensals. We performed long-read sequencing using Oxford Nanopore technology and used this together with pre-existing Illumina sequences to *de novo* assemble these into high quality genomes with improved contiguity and quality. These genomes now provide a solid platform for detailed, mechanistic understanding of microbiome assembly, dynamics and evolution in plants.

## Background & Summary

In recent decades the importance of the plant microbiota has been well established, with burgeoning research efforts aiming to extend understanding of plant microbiomes (‘the plant’s second genome’) for improvement of plant health, survival, and growth^1,2^. The soil environment in particular contains some of the greatest biological diversity known, and the rhizosphere (influenced by plant root exudates) contains up to 10^11^ microbes per gram of plant root^1^. However, this also means that within the rhizosphere, microbes exist in incredibly complex and dynamic communities^2,3^. To help disentangle the complexity of host-microbe and microbe-microbe interactions in these settings, reductionist experimental approaches such as the use of Synthetic Communities (SynComs) are increasingly being utilized^4–7^.

SynComs consist of previously isolated bacteria, spanning diverse bacterial taxa, that covers the natural taxonomic diversity and composition from a particular environment^4,8^. These bacteria can be cultured under controlled conditions, to investigate characteristics and dynamics within the bacterial community and/or community-host interactions^9^. The application of bacterial SynComs across various research areas has yielded interesting discoveries in, for example, bioremediation^10,11^, disease treatment^12–14^, plant growth promotion^15,16^, or plant biotic^17,18^, and abiotic stress mitigation^19^.

*Lotus japonicus* is a model legume that is widely used for understanding rhizobia-legume interactions^20,21^. The *Lotus japonicus* culture collection^22^, also called *Lj*-SPHERE, was established by isolation of commensal bacteria from the root compartment of *L. japonicus* grown in Cologne agricultural soil. The isolates were whole genome sequenced and assembled using Illumina short reads, followed by taxonomic identification using amplicon sequencing of the 16S ribosomal gene marker^22^. The previous assemblies existed only as low-quality draft genomes to enable 16S rDNA amplicon-based abundance profiling of the community.

Here, we aimed to improve the *Lj*-SPHERE low-quality drafts to obtain contiguous and annotated high-quality genomes as reference material for future studies. For that, we focused on a subset of 177 taxonomically diverse genomes that were previously determined as unique through a one-to-one whole genome nucleotide similarity comparison. Genomes with similarity values of 99.99% or higher were clustered together, and within each cluster (n=41), one genome was chosen as a representative. The isolates, for the 177 genomes, were re- grown and processed for DNA extraction, library preparation, and long-read sequencing using the Oxford Nanopore Technology (ONT) MinION sequencing platform.

We obtained ONT long-read sequences for all the targeted isolates, allowing us to re- assemble these genomes using a hybrid assembly approach by combining the original Illumina reads with newly produced long reads (Figure 1a). Inspection of the available Illumina sequences revealed that for four isolates novel Illumina sequencing was required to reach the necessary quality for the input data. The assembly pipeline (Figure 1a & 1b) provided us with 152 high-quality annotated bacterial genomes, where a large number of genomes were generated automatically (n=148) and only a small number of genomes required manual curation for completion (LjRoot30, LjRoot166, LjRoot111 and LjRoot112A) (Figure 1c). To ensure high quality assemblies, we implemented genome validation thresholds based on the National Center for Biotechnology Information (NCBI) Prokaryotic Genome Annotation Standards. These thresholds were defined as minimum completeness of 80%, less than 5% genomic contamination and, at least one complete copy of the required ribosomal genes (5S, 16S and 23S). The overall statistics performed on the final genomes (N50 and L50), revealed that 152 genomes were successfully assembled with good contiguity and high quality (Figure 2; Supplementary Table 1).

**Figure 1:**
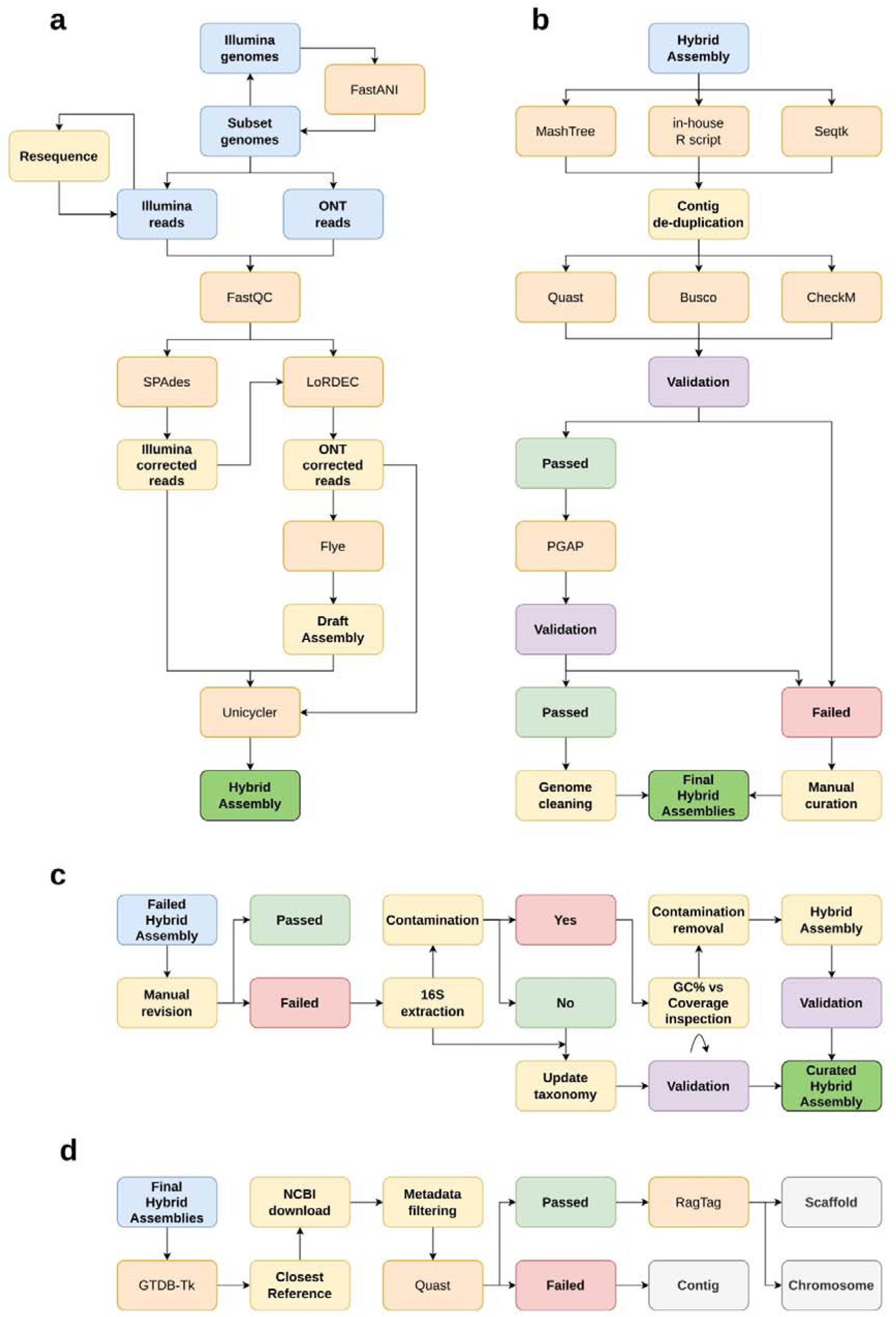
Schematic overview of the hybrid assembly pipeline and downstream workflow. a) Hybrid assembly pipeline starting from the input sequencing data and finishing with a hybrid assembly draft. b) Computational workflow to validate and obtain a final hybrid assembly. c) Manual curation workflow including 16S taxonomic identification and reclassification. d) Genome scaffolding workflow. Across panels, the colour pattern represents: input (blue), software (orange), computational step (yellow), validation (purple), output (bright green), category (light grey), and passed/failed (green/red).

**Figure 2:**
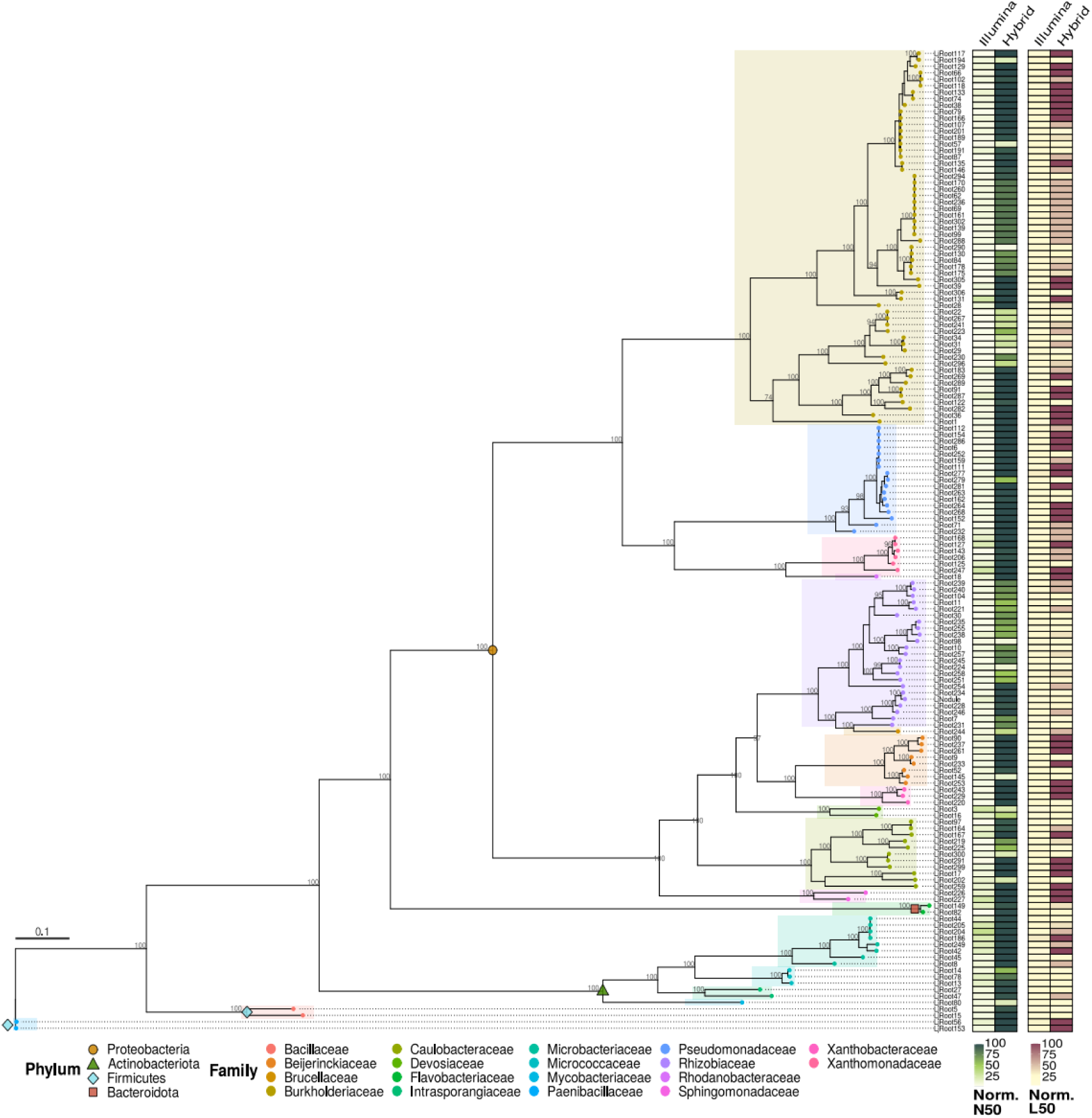
A maximum likelihood phylogenetic tree was constructed using concatenated alignments of 20 validated bacterial core genes (VBCG) across the 152 improved *Lj-*SPHERE genomes. The numbers at branch nodes represent 100 bootstrap replicates. The bar represents the number of substitutions per nucleotide site. Phylum is represented by four different symbols before the branch nodes. Branches are grouped as taxonomic families by coloured rectangles and tree tips. The first heatmap (green) displays the percentage of genome length covered by the N50 metric given the total length. The second heatmap (yellow/maroon) displays the percentage of genome contigs by the L50 given the total number of contigs. Both heatmaps contain two columns, the first is Illumina *Lj-*SPHERE genomes and the second is the hybrid assembled set of improved genomes. For both scales (Norm. N50 & L50), darker values determine higher quality metrics.

The hybrid assembled genomes were taxonomically classified using GTDB-tk^23^, that was utilized to provide a revised taxonomic classification and an estimate of the closest available references supported by nucleotide-wise comparisons. Inspection of the distribution of taxonomic identification across the 152 genomes revealed a majority of Proteobacteria (n=125), followed by other phyla, such as Actinobacteriota, Bacteroidota, and Firmicutes (Figure 2). At the family level, we identified 18 distinct taxa and a predominance of Burkholderiaceae (n=58), Rhizobiaceae (n=23), Pseudomonadaceae (n=17), and Caulobacteraceae (n=11). Furthermore, the results revealed a large overlap between the previous 16S ribosomal gene marker and the newly produced taxonomy at the genus level. A small number of genomes underwent a shift from their previous identified genus (n=27), although the majority of them were subsequently labeled as a sister genus (n=15). A considerable number of genomes were identified reliably at the species level (n=58), vastly improving the taxonomic assignment available for the *Lj-*SPHERE collection.

To ensure the highest quality genomes, we identified the closest available references from NCBI based on taxonomic classification and used these to scaffold the hybrid-assembled genomes (Figure 1d). This revealed that out of 152 genomes, only 35 of these could use the closest reference(s) from NCBI as a scaffolding reference. The remainder of our genomes had a lower fragmentation rate and higher overall genome quality compared to the relevant NCBI references, therefore the NCBI references could not be used. This demonstrates the high quality of the reassembled *Lj-*SPHERE genomes reported here, which are publicly available for the scientific community at the European Nucleotide Archive (ENA).

Overall, we have produced 152 high-quality annotated genomes of commensal bacteria from the roots of *L. japonicus,* substantially improving this resource for comprehensive bioinformatic analyses, including metagenomic analysis, gene cluster identification, comparative genomics, and genome mining. This also creates the necessary foundation for future experimental studies such as those focused on bacterial evolution and whole genome mutagenesis, where a high-quality genome is required.

## Materials & Methods

### Acquisition of previous genomic data and Illumina sequencing reads

The genome assemblies for the *Lj-*SPHERE were downloaded from the *At*-SPHERE website (https://www.at-sphere.com/). The 16S ribosomal gene taxonomic identification of the downloaded genomes was obtained from the Supplementary Data previously published^22^. The corresponding pre-existing Illumina reads, in fastq format, were downloaded from the European Nucleotide Archive or ENA (Project: PRJEB37696). Inspection of the paired end reads revealed four isolates (LjRoot112A, LjRoot204, LjRoot205 and LjRoot206) that contained an inadequate number or quality of Illumina reads, therefore were re-sequenced. The extracted DNA (described below) was sent to Novogene (Munich, Germany) for library preparation and paired end Illumina sequencing (150 bp; Novaseq 6000).

### Subset of Illumina genomes

Pairwise comparisons of nucleotide similarity were computed between the Illumina genomes using FastANI (v1.33)^24^. The genomes with similarity values of 99.99% ANI or higher were clustered together whereas the genome with similarity values below this threshold were considered unique entities. A representative candidate for each genome cluster was selected based on the optimal balance between the contig number and the genome length, while prioritizing the selection of largely contiguous genomes. The Illumina reads of the non- selected cluster candidates were merged to increase assembly coverage for the representative isolate. The resulting list, consisting of the cluster representative isolates along with the unique isolates, were selected for long read sequencing.

### High Molecular Weight DNA extraction

The subset of selected genomes of previously isolated, purified and stored *L. japonicus* strains^22^ were streaked from glycerol stocks onto 0.3% Tryptic Soy Agar (TSB) plates. Pellets were prepared by re-suspending a full loop of bacterial biomass into 500 μl of sterile distilled water and centrifuging at 9000g for 2 min. High Molecular Weight (HMW) DNA was extracted from the pellets as detailed previously^25^, with modifications for the lysis step. Briefly, pellets were resuspended in 500 μl lysis buffer A (10 mg/ml lysozyme, 100 mM Tris-HCl pH 8.0, 50 mM EDTA pH 8.0) and incubated for 30 min at 37°C. Then, 300 μl of lysis buffer B (1.3 M NaCl, 100 mM Tris-HCl pH 8.0, 50 mM EDTA pH 8.0 and 4% SDS), 4 μl of RNase A (20 mg/ml), and 8 μl of proteinase L (20 mg/ml) were added to the mix and incubated for 20 min at 60°C. Protein and polysaccharide precipitation was achieved by adding 0.3x volumes of 5 M potassium acetate and incubating at 4°C for 10 min. The mix was centrifuged at 12,000g for 20 min and the supernatant was transferred to a new low-bind Eppendorf tube. DNA was recovered by adding 1 ml of binding buffer (20% PEG 8000 and 3 M NaCl) and 100 μl of AMPure XP beads by using a magnetic rack. Finally, DNA was washed twice with 70% ethanol, resuspended in 10 mM Tris-HCl pH 8.0, and fluorometrically quantified using Qubit™ dsDNA HS kit. Samples were stored at 4°C until use.

### ONT platform sequencing

Diluted DNA samples (50 ng) were used as input for library preparation. Genomic DNA libraries were prepared using the Rapid Barcoding Kit 96 V14 (SQK-RBK114.96) and sequenced on R9.4.1 flow cells using the ONT MinION sequencing device^26^. Each round was processed and monitored using the ONT MinKNOW software (v21.11.9).

### Genome reconstruction using a hybrid assembly approach

ONT fast5 files were base called and demultiplexed into fastq files using Guppy (v6.1.1). The ONT reads (FASTQ) together with the Illumina reads (FASTQ), were supplied to an in-house pipeline to perform the hybrid assembly and subsequent annotation of the resulting bacterial genomes. Our pipeline, *HA.py* (v1.2), implemented using gwf (v1.7.2) and partly adapted from a previously published pipeline^27^, started with quality control (QC) using NanoPlot (v1.36.1)^28^ and FastQC (v0.11.9) for the ONT and Illumina reads, respectively. The QC results were assessed before proceeding. Illumina reads that passed the QC were corrected for sequencing errors using SPAdes (v3.15.4)^29^, then passed to LoRDEC (v0.9)^30^ to correct sequencing errors in the ONT reads. Corrected long reads were passed to Flye (v2.9-b1768)^31^ to generate a draft assembly using only the ONT reads. The three sources of information, corrected short reads, corrected long reads, and ONT draft assembly, were used as input to perform the hybrid assembly using unicycler (v0.5.0)^32^. The resulting genome was cleaned of duplicated contigs, or contigs within larger contigs, using Mashtree (v1.2.0)^33^, an in-house R script, and seqtk (v1.3-r106).

Genome quality was checked using Quast (v5.0.2)^34^, BUSCO (v5.3.2)^35^, and CheckM (v1.2.0)^36^. The BUSCO validation required complete and single genes to be above 90%, and duplicated, fragmented, and missing genes below 5%. Moreover, CheckM validation required completeness above 90% and contamination below 5%. If the hybrid assembly passed the validations, it was then annotated using the Prokaryotic Genome Annotation Pipeline, or PGAP, from the NCBI^37^.

The PGAP annotation was used to validate the quality of the genome. Genomes were deemed successful if they contained the following: a gene ratio above 80%, less than 20% pseudogenes, at least 20 tRNAs, and at least one copy of the 5S, 16S, and 23S rRNA subunits. For this validation the identified 5S, 16S, and 23S rRNA sequence lengths were allowed ≤10% deviation compared to the *Escherichia coli* rRNA subunits (5S: 120 bp, 16S: 1542bp, and 23S: 2904 bp). Finally, contigs containing no genes or only pseudogenes were considered spurious and therefore discarded from the genome and the annotation.

### Manual curation

Manual curation of unsuccessful genome assemblies was performed depending on the cause of the failure. First, when CheckM reported contamination greater than 5%, we used an auxiliary pipeline to search for contamination within the original Illumina genomes and, in parallel, within the ONT-based draft assemblies (an intermediate product from the hybrid assembly pipeline). The auxiliary pipeline identified the 16S ribosomal gene using Blastn (v2.12.0+)^38^, removed sequence duplication using seqkit (v2.2.0)^39^, then extracted the V3-V4 and V5-V7 subunits using QIIME2 (v2022.11.1)^40^. The SILVA database (v138)^41^ was downloaded and converted into 16S, V3-V4 and V5-V7 QIIME2 format databases and used for taxonomical classification. The taxonomic results were used to identify sources of contamination originating from the Illumina and/or ONT input reads.

When BUSCO encountered discrepancies between the *Lj-*SPHERE 16S ribosomal gene and the hybrid assembly taxonomies, we utilized the aforementioned 16S, V3-V4 and V5-V7 taxonomic identifications, checked for concordance with the CheckM identification and then updated the *Lj*-SPHERE taxonomy accordingly. The hybrid assembly was then re-run using the corrected taxonomic information, enabling successful validation.

### Taxonomic classification and phylogeny

The resulting hybrid assembled genomes were taxonomically classified using the GTDB-Tk toolkit (v2.1.1)^23^, and the resulting taxonomy was compared with the updated 16S taxonomic identification (Manually curated SILVA database) to evaluate the degree of congruence and account for potential discrepancies. GTDB-Tk was then utilized to determine the taxonomy of the genomes to species level, or if not possible, to unidentified species level. A phylogenetic tree was constructed based on the 20 validated bacterial cores genes (VBCG) determined and utilised by the VCBG pipeline (v1.3)^42^.These sequences were then fed into RAxML (v8.2.12)^43^ to construct a maximum likelihood phylogenetic tree with 100 bootstrap replicates. The resulting phylogeny was plotted using the R package ggtree (v3.10.0).

### Genome categories for ENA upload

GTDB-Tk toolkit^23^ taxonomic identification results were used to identify the closest genome references. The software placed each genome on the bacteria and archaea tree of life, reporting the closest and the second closest reference, and a clade comprised by comparable references. Hybrid assembled genomes consisting of more than one contig were considered for scaffolding. The appropriate scaffold reference was selected sequentially along the closest, second closest or clade references, considering for the latter only references with ANI values above 80% similarity. Once selected, the references were downloaded from GenBank or RefSeq and their quality (N50, L50, number of contigs, and genome length) was assessed using the available metadata and Quast (v5.0.2)^34^. Among the downloaded references, only the genomes with better quality metrics than our hybrid assembled genomes were used for genome scaffolding. Hybrid assembled genomes were scaffolded using RagTag (v2.1.0)^44^. Finally, we manually inspected the resulting scaffolded genomes and determined which ENA category (contig, scaffold, or chromosome) was most appropriate for data submission. Genome assemblies with one contig were directly included in the chromosome category, whereas genomes that were not scaffolded were defined as contig category.

### Data records

The improved set of genomes, their corresponding ONT reads, and the Illumina reads for the four Illumina re-sequenced isolates are available at the European Nucleotide Archive (ENA) under the project PRJEB75483.

### Technical validation

#### Bacterial re-growth, DNA extraction and library preparation

Isolates of *Lj-*SPHERE were cultured in sterile conditions on 0.3% TSB medium. They were checked for purity based on single colony morphology and handled using aseptic techniques. DNA extractions were carried out in a specialised laboratory environment and all equipment was sterilised before use.

#### Genome assembly

The technical validation of the genomes consisted of:

1. Quality control of raw reads.
2. Taxonomy verification using multiple software.
3. Annotation validation following the NCBI Prokaryotic Genome Annotation Standards (https://www.ncbi.nlm.nih.gov/refseq/annotation_prok/standards/).
4. Assembly redundancy and spurious contig removal.
5. Genome scaffolding, when applicable.

### Code availability

The in-house pipeline, HA.py (v1.2), the scaffolding pipeline, scaffolding.py (v1.0), and the auxiliary pipeline, extract16Sregion.py (v1.0), including all the scripts and Anaconda environments used for the generation of this dataset, can be found on GitHub (https://github.com/adriangeerre/HybridAssembly).

## Acknowledgments

This work was supported by the project Molecular Mechanisms and Dynamics of Plant– Microbe Interactions at the Root–Soil Interface (InRoot), supported by the Novo Nordisk Foundation (grant number NNF19SA0059362) (https://ccrp.vcl.ncsu.edu/content/inroot).

## Author contributions

S.F conceived and designed the study. E.V.R performed the microbiology, DNA extraction, and sequencing library preparation. S.F and E.V.R performed the ONT sequencing. A.G-R designed and executed the computational methodology, including data collection, processing, hybrid assembly, annotation, and verification of the genomes. A.G-R wrote the first draft of the manuscript with contributions from E.V.R. S.R. and S.F supervised the study and revised the manuscript. All authors contributed to reviewing the draft versions of the manuscript. S.R provided the funding and resources to conduct the study.

## Competing interests

The authors declare no competing interests.

## Additional information

### Supplementary information

Supplementary Table 1: Assembly statistics of Illumina and Hybrid assembled strains

## Notes

https://www.ebi.ac.uk/ena/browser/view/PRJEB75483

### Competing Interest Statement

The authors have declared no competing interest.

### Summary of Updates

We revised and improved terminology used in the original submission.

